# Biogeographic Ancestry and Socioeconomic Outcomes in the Americas: A Meta-Analysis

**DOI:** 10.1101/055681

**Authors:** Emil O. W. Kirkegaard, Mingrui Wang, John Fuerst

## Abstract

Narrative reports suggest that socioeconomic status (SES) is associated with biogeographic ancestry (BGA) in the Americas. If so, SES potentially acts as a confound that needs to be taken into account when evaluating the relation between medical outcomes and BGA. To explore how systematic BGA-SES associations are, a meta-analysis of American studies was conducted. 40 studies were identified, yielding a total of 64 independent samples with directions of associations, including 48 independent samples with effect sizes. An analysis of association directions found a high degree of consistency. The square root *n*-weighted directions were 0.83 (*K* =36), -0.81 (*K* = 41) and -0.82 (*K* = 39) for European, Amerindian and African BGA, respectively. An analysis of effect size magnitudes found that European BGA was positively associated with SES, with a meta-analytic effect size of *r* = .18 [95% CI: .13 to .24, *K* = 28, *n* = 35,476.5], while both Amerindian and African BGA were negatively associated with SES, having meta-analytic effect sizes of -.14 [-.18 to -.10, *K* = 31, *n* = 28,937.5] and -.11 [-.15 to -.07, *K* = 28, *n* = 32,710.5], respectively. There was considerable cross-sample variation in effect sizes (mean I^2^ = 92%), but the sample size was not enough for performing credible moderator analysis. Implications for future studies are discussed.

---

Admixture analysis is a potent tool for the exploration of the etiology of traits and trait differences in admixed populations. Admixture mapping is a form of admixture analysis that allows for the detection of specific disease- and trait-associated genes (Shriner, 2013; Winkler, Nelson & Smith, 2010). When biogeographic ancestry (BGA) groups differ in the frequency of disease- or trait-causing genetic variants, in admixed populations the phenotype of interest will be correlated with BGA in genomic regions near the causal genetic variants, allowing for the identification of associated loci. Such associations will occur *within* self-identified race/ethnicity (SIRE) groups. When the genetic architecture of a trait is complex, with hundreds of loci assumed to contribute to the phenotype, an analysis of global admixture is an appropriate first step. This strategy seeks to identify associations of global admixture proportions with the measured trait, without attempting to identify individual loci. Unlike admixture mapping, which requires sample sizes of many thousands, global admixture analysis can produce meaningful results with smaller sample sizes.

As used here, *population* refers to geographically delineated population. In contrast, *BGA* refers to genetic ancestry as defined by the presence of ancestrally informative molecular markers in a person’s genome, which allow the calculation of an individual’s resemblance to the average genotypes in the ancestral populations. These ancestral populations have also been called clusters or ancestral groups (e.g., Frudakis, 2010; Frudakis & Shriver, 2003; Shriver & Kittles, 2004). With respect to studies of American populations, the ancestral populations are typically Europeans, Sub-Saharan Africans, and Amerindians (Salzano & Sans, 2014).

Socioeconomic status (SES) inequalities between and within SIRE groups can lead to spurious associations of BGA with medical outcomes when both are associated with SES (Marden et al., 2016). Thus, genetic traits associated with SES whose allele frequencies differ among ancestral groups can be misidentified as being associated with a specific medical outcome in admixture mapping. For this reason, controls for SES are not infrequently incorporated into such analyses. Narrative reports (e.g., González Burchard et al., 2005) have suggested that SES covaries with admixture such that, relative to European BGA, Amerindian and African BGA are associated with lower SES. If this is typically the case, it would be prudent for researchers to include reliable measures of SES as covariates in analyses to provide lower-bound estimates of the BGA-medical outcome associations. Moreover, it would be advisable to investigate the causal pathways mediating the BGA-SES associations, to identify possible unobserved non-genetic mediators of BGA-medical outcomes associations. However, no meta-analysis has been conducted to date to establish whether SES outcomes are associated with BGA in any consistent way.

## 1. Methods

### 1.1. Collection of studies and data exclusion

Papers were sought which reported quantitative or qualitative associations between European, African, or Amerindian BGA and socioeconomic outcomes. In 2014/15, a literature search was conducted using the PubMed and BIOSIS previews databases. Searches such as the following were used: ‘(admixture) AND (socioeconomic or education or income or SES or poverty) AND (African or European or Amerindian)’. A search, limited to the years 2003 to early 2016, was also conducted using Google Scholar. For the latter, phrases such as ‘(ses, OR income, OR education) AND (genomic OR biogeographical, OR ancestry, OR admixture) AND (African, OR European, OR Amerindian)’ were used. The Google Scholar search yielded over twenty thousand hits in descending order of relevance to the search terms; the first 1,500 abstracts were skimmed and approximately 250 papers were identified as potential sources and read. The references of the identified papers, from the PubMed, BIOSIS, and Google Scholar searches, were inspected for additional sources. In total, we identified 54 papers (51 from the database search and three from the reference search). Of these, six used redundant samples and thus were excluded. For eight of the papers, while SES was a covariate in an admixture analysis, no usable information was recoverable and authors did not reply to emails sent to their listed emails. The remaining 40 papers either contained directions of associations or these were provided by the authors when contacted; for 27 of the 40 papers, either effect sizes (or metrics that could be converted into effect sizes) were reported or provided by the authors. Figure 1 shows a flowchart of the procedure.

**Figure 1.**
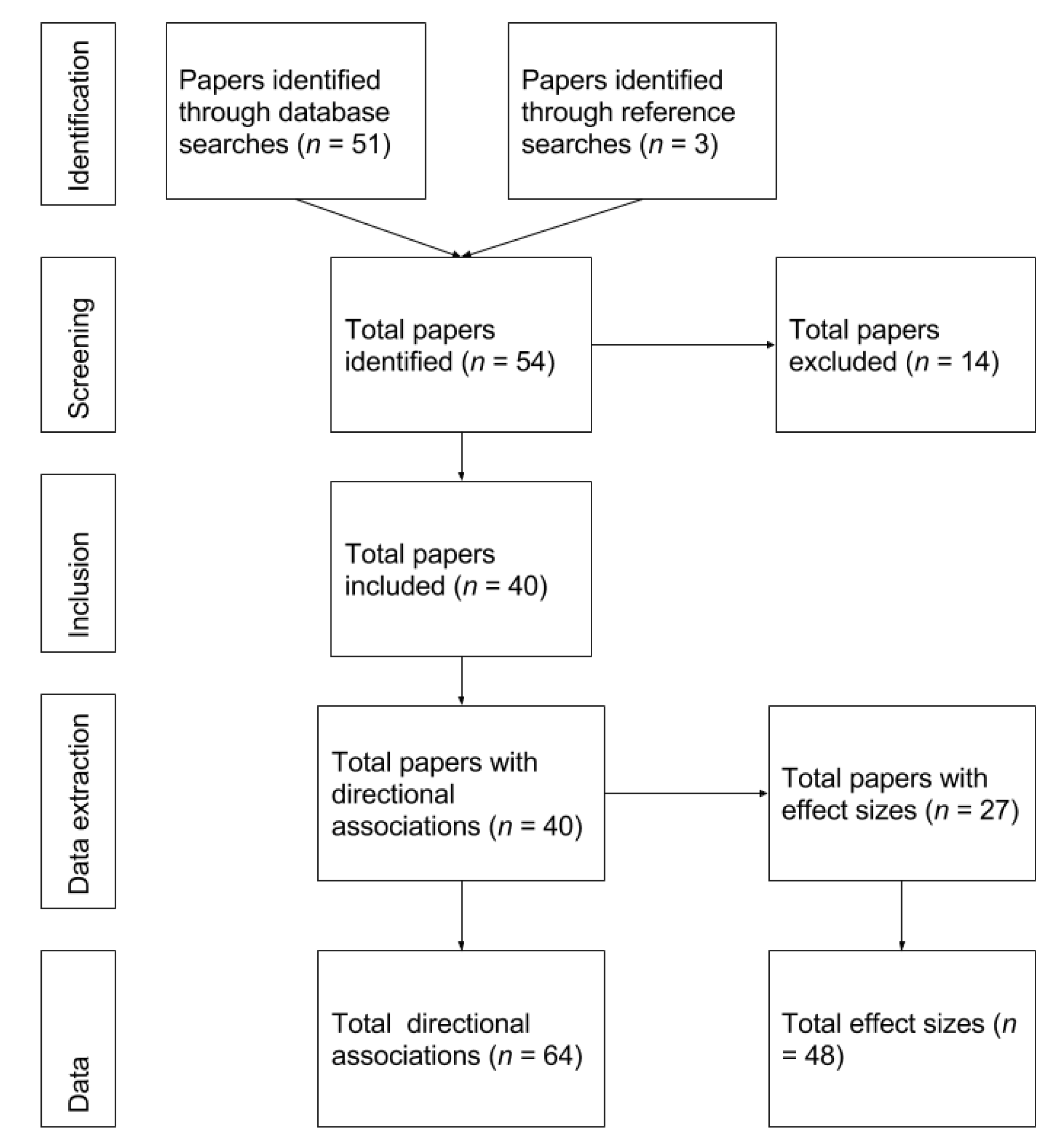
Flowchart of study inclusion.

The identified studies and reasons for exclusions are listed in Supplementary File 2. Relevant information from each study and sample was recorded and is reported in Supplementary File 1. With respect to coding, two of this paper’s authors reviewed studies and reached agreement on ambiguous cases. Many studies did not report effect sizes or statistics convertible to effect sizes; some studies failed to clearly report association directions. For studies that did not report effect sizes or directions of associations, authors were contacted. Contact dates, details, and notes are reported in Supplementary File 1 along with annotated replies when forthcoming. Seventy-three percent of our requests for data elicited a reply. For 38 percent of requests the authors provided results not reported in the original papers.

Most papers were published in the last few years, with the median year of publication being 2015 (range: 1988 to 2016). Many studies had medical themes and often included both case and control samples. Table 1 shows the independent samples broken down by sample type. Samples were coded as ‘medical’ versus ‘non-medical’ based on the authors’ research goal. As a result, for some samples, this classification was somewhat arbitrary. For example, Zou et al. (2015) explored genetic assortative mating using the Genes-Environments and Admixture in Latino Americans (GALA-II) community survey. Since the outcome of interest was ‘non-medical’, the samples were classified as ‘non-medical’. However, if the same authors had instead explored the relation between BGA and hypertension, using the same samples, and if results were undifferentiated by clinical conditions, the samples would have been classified as ‘case and control’ (meaning samples which include both cases and controls).

**Table 1.** Independent samples by sample type

| Sample type | Direction of associations available | Effect sizes available |
| --- | --- | --- |
| Case | 10 | 8 |
| Case and control | 24 | 19 |
| Control | 8 | 4 |
| Non-medical | 22 | 17 |
| Total | 64 | 48 |

For the meta-analysis, we report results based on associations for which the relevant BGA proportions are above 5%. For comparison, we also note the results with low-admixture samples included. Associations in cases of low admixture were excluded because when means and thus variances of admixture are low, effect sizes will be attenuated. Regarding unreliability, one concern is that the present meta-analysis is based on correlation coefficients. As such, the statistical effects of other BGA components are not held constant and covariance between components can confound associations, leading to spurious results. Possible confounding is more of a concern when admixture fractions are low. With this exclusion, the total number of independent samples (64 for directions of associations and 48 for effect sizes) remained the same, but the number of samples when decomposed by specific BGAs was reduced. The number of samples is shown in Table 2.

**Table 2.** Number of independent associations by BGA without (w/o) and with (w) low-admixture samples.

|  | Independent samples | Associations (African) | Associations (Amerindian) | Associations (European) |
| --- | --- | --- | --- | --- |
| Directional associations (w/o low admix) | 64 | 39 | 41 | 36 |
| Directional associations (w low admix) | 64 | 43 | 43 | 36 |
| Effect sizes (w/o low admix) | 48 | 28 | 31 | 28 |
| Effect sizes (w low admix) | 48 | 31 | 33 | 28 |

Only a subset of the independent samples reported associations for all three BGAs. As a result, the number of associations for a particular BGA is less than the number of independent samples. When studies reported associations between outcomes and one or two BGA components, one could attempt to estimate association directions and effect sizes for the remaining BGAs since the three BGAs are directly related to one another, with ancestries adding up to unity. For instance, Bonilla (2015, extra information provided by the author) reported that European and Amerindian BGA correlated at -.82 and -.85 in two samples (*ns* = 148, 164). The correlations between European BGA and SES were .10 and .14, respectively; thus, one might infer that the Amerindian ⨯ SES correlations were -.10 and -.14. In this case, however, they were -.01 and -.13; the departure from expectation resulted from the association between African BGA and SES. To avoid estimation error, unreported associations and effect sizes were not estimated.

### 1.2. Descriptive statistics of studies

In some instances, associations with multiple outcome measures were reported for a single sample and the same BGA. Given this situation, treating BGA ⨯ outcome associations as independent data points would lead to double counting. Two recent meta-analyses encountered an analogous problem (Tucker-Drob & Bates, 2015; Tate & McDaniel, 2008). The method employed by the first was to use a complex weighting approach to avoid double counting, while Tate and McDaniel (2008) used a simpler approach of averaging results within samples before aggregating. An approach similar to the second was implemented in the present study. Median values, which are more robust to outliers than are means, were taken across outcome values for the same BGA within each sample before meta-analyzing the sample associations. Different classes of outcomes (e.g., education vs. income) were not weighted differently. As Tate and McDaniel (2008) point out, this method slightly throws off the standard errors, but this problem was judged to be minor. Henceforth, all results are reported for independent samples (with the exception of Table 3), with BGA ⨯ outcome associations having been first averaged within each sample.

**Table 3.** Broad outcomes (specific outcomes) for all BGA ⨯ outcome associations with/without low admixture excluded

| Broad outcome (specific outcomes) | Frequency | Broad outcome (specific outcomes) | Frequency |
| --- | --- | --- | --- |
| Education (education, years of schooling) | 69/73 | Parental SES (parental Hollingshead index, running water, had a car) | 6/6 |
| Income (income, wealth index, household income per capita) | 49/53 | Parental education | 2/2 |
| SES (SES, SES + education, occupation, general factor of SES, Hollingshead index, Yost index, home property band, household asset index) | 34/35 | Parental income | 1/1 |
| Neighborhood SES | 18/19 | Total | 179/189 |

For the 64 independent samples, there were 179 BGA ⨯ outcome associations, since some samples included data for multiple BGAs and multiple SES indicators. Studies reported a large variety of specific outcome variables (Table 3). As shown in Supplementary File 1, these variables were coded into broad categories. Supplementary Table 1 shows the breakdown of all outcomes by broad category. Most measures were of individual education, income or occupation. A few BGA ⨯ SES associations were based on the socioeconomic level of the participant’s neighborhood. Depending on the model of the proposed covariance, this could be a questionable index. For example, a recent study based on the UK Biobank (N ≈ 112k) found a modest phenotypic correlation between individual and neighborhood-level SES (.24) but a strong genetic one (.87) (Hill et al., 2016). While neighborhood-level measures are included in the meta-analysis, it is advisable that future investigators conduct moderator analyses to estimate the impact of using neighborhood versus individual indicators. Additionally, several studies reported parental SES. Because most children are the biological offspring of both of their parents, their admixture will index the average of their parents, and these results can then be seen as showing the correlation between the parents’ SES and the parents’ BGA. Roughly sixty percent of the 64 independent samples came from the US. Studies often divided their samples into SIRE groups. Tables 4a and 4b show a breakdown by country and SIRE group.

**Table 4a.** Number of independent samples and sample sizes for directional association analysis by SIRE group and country. Low admixture excluded.

| SIRE group | K | N | SIRE group | K | N | SIRE group | K | N |
| --- | --- | --- | --- | --- | --- | --- | --- | --- |
| African American (US) | 10 | 20581 | Multi-ethnic (Chile) | 2 | 1967 | African-descent (T&T) | 1 | 107 |
| Hispanic (US) | 10 | 9983 | Multi-ethnic (Mexico) | 2 | 1991 | Multi-ethnic (Argentina) | 1 | 220 |
| Mexican (US) | 5 | 1050 | Multi-ethnic (US) | 2 | 2031 | Multi-ethnic (Colombia) | 1 | 2092 |
| Multi-ethnic (Brazil) | 6 | 6696 | Mestizo (Chile) | 2 | 2381 | Not stated (Costa Rica) | 1 | 1998 |
| Puerto Rican (PR) | 7 | 2368 | Native American (US) | 2 | 741 | Puerto Rican (US) | 1 | 122 |
| Not stated (Mexico) | 3 | 1831 | Not stated (Uruguay) | 2 | 312 | White American (US) | 1 | 73 |
| Multi-ethnic (Peru) | 3 | 1979 | Not stated (Colombia) | 2 | 1971 | Total | 64 |  |

*Note:* N = participant sample size; K = number of independent samples. PR = Puerto Rico; T&T = Trinidad and Tobago.

**Table 4b.** Number of independent samples and sample sizes for effect size analysis by SIRE group and country. Low admixture excluded.

| SIRE group | K | N | SIRE group | K | N | SIRE group | K | N |
| --- | --- | --- | --- | --- | --- | --- | --- | --- |
| African American (US) | 7 | 13589 | Multi-ethnic (Chile) | 2 | 1967 | African-descent (T&T) | 0 |  |
| Hispanic (US) | 6 | 8041 | Multi-ethnic (Mexico) | 2 | 1991 | Multi-ethnic (Argentina) | 1 | 220 |
| Mexican (US) | 5 | 1050 | Multi-ethnic (US) | 2 | 2031 | Multi-ethnic (Colombia) | 1 | 2092 |
| Multi-ethnic (Brazil) | 5 | 6518.5 | Mestizo (Chile) | 0 |  | Not stated (Costa Rica) | 0 |  |
| Puerto Rican (PR) | 3 | 1943 | Native American (US) | 2 | 741 | Puerto Rican (US) | 1 | 122 |
| Not stated (Mexico) | 3 | 1831 | Not stated (Uruguay) | 2 | 312 | White American (US) | 1 | 73 |
| Multi-ethnic (Peru) | 3 | 1979 | Not stated (Colombia) | 2 | 1971 | Total | 48 |  |

### 1.3. Directions of associations

As shown in Table 2, directions of associations were more frequently available than were actual effect sizes. Association directions can provide some indication as to whether the findings are in line with a null hypothesis. Other meta-analyses have included analyses of association directions (e.g., Van der Meer & Tolsma, 2014).

### 1.4. Statistical analysis

For each sample, directions (negative, null, positive) were assigned to associations. With respect to this assignment, neither *p*-values nor the magnitude of effects were taken into account. Thus, for example, the correlation between African BGA and education for Bonilla et al.’s (2015) sample 1 was -.16 (*n* = 164, *p* = .08), while the correlation between African BGA and education for Bonilla et al.’s (2015) sample 2 was .04 (*n* = 148, *p* = .59); the first association was coded as ‘negative’ and the second as ‘positive’ despite the difference in effect size. Those samples coded as ‘null’ were ones where the effect size was zero. After directions were assigned, they were coded as negative = –1, null = 0, and positive = 1). If a sample had consistent directions (across multiple SES indicators), it received a score of either -1 or 1. If a sample had mixed results, it received an intermediate score equivalent to the mean of the different associations (e.g., a sample reporting three positive and one negative associations received a score of (3-1)/4=0.5.) The results are reported unweighted; weighted by the square root of the sample sizes; and weighted by the sample sizes. We consider square root *n*-weighted results to be preferable as they strike a good balance between taking into account the effect of sample size and not obscuring results from smaller samples; other researchers have employed this weighting strategy (for example, Kan et al., 2013).

## 2. Results

### 2.1. Directions of associations

The aggregated within-sample directions are shown in Table 5. In general, the directions are positive for European BGA and negative for the other two ancestries. With respect to African and Amerindian BGA, the weighted results are substantially stronger when low-admixture samples are excluded. There were no samples with low European BGA.

**Table 5.** Directions of associations by BGA

| BGA | N | K | Mean direction<br>(unweighted) | Mean direction<br>(SQRT(n)-<br>weighted) | Mean direction<br>(n-weighted) |
| --- | --- | --- | --- | --- | --- |
| African (w/o low admix) | 40931.5 | 39 | -0.67 | -0.82 | -0.94 |
| African (w low admix) | 47742.5 | 43 | -0.75 | -0.64 | -0.56 |
| Amerindian (w/o low admix) | 35577.5 | 41 | -0.76 | -0.81 | -0.85 |
| Amerindian (w low admix) | 47107.5 | 43 | -0.72 | -0.65 | -0.41 |
| European (w/o low admix) | 39548.5 | 36 | 0.72 | 0.83 | 0.89 |
| European (w low admix) | 39548.5 | 36 | 0.72 | 0.83 | 0.89 |

### 2.2. Effect sizes

For 92% of the samples with effect sizes, Pearson correlations or other effect sizes that could be directly converted into these (i.e., r^2^, Spearman’s rho, beta coefficients, and odds ratio) were available. For other samples, effect sizes needed to be computed from t-tests (N = 1), *F*-tests (N = 4), and frequency tables (N = 5). Conversions were made using the formulas and methods noted in Supplementary File 1. To provide an overview of the effect sizes, a density histogram plot was made of the correlations for each BGA component. Additional statistics for the effect sizes were also computed. For the meta-analysis, because samples were from different populations in different countries, a random effects model was appropriate (Schmidt & Hunter, 2014). We conducted the analysis using the *metafor* package for R(Viechtbauer, 2015).

The studies varied substantially in sample size. Figure 2 shows the distribution of the sample sizes for the effect size analysis with low-admixture samples excluded. The density histogram plot of the effect sizes is shown in Figure 3. Descriptive statistics for the effect sizes with and without low admixture samples excluded are shown in Table 6.

**Figure 2.**
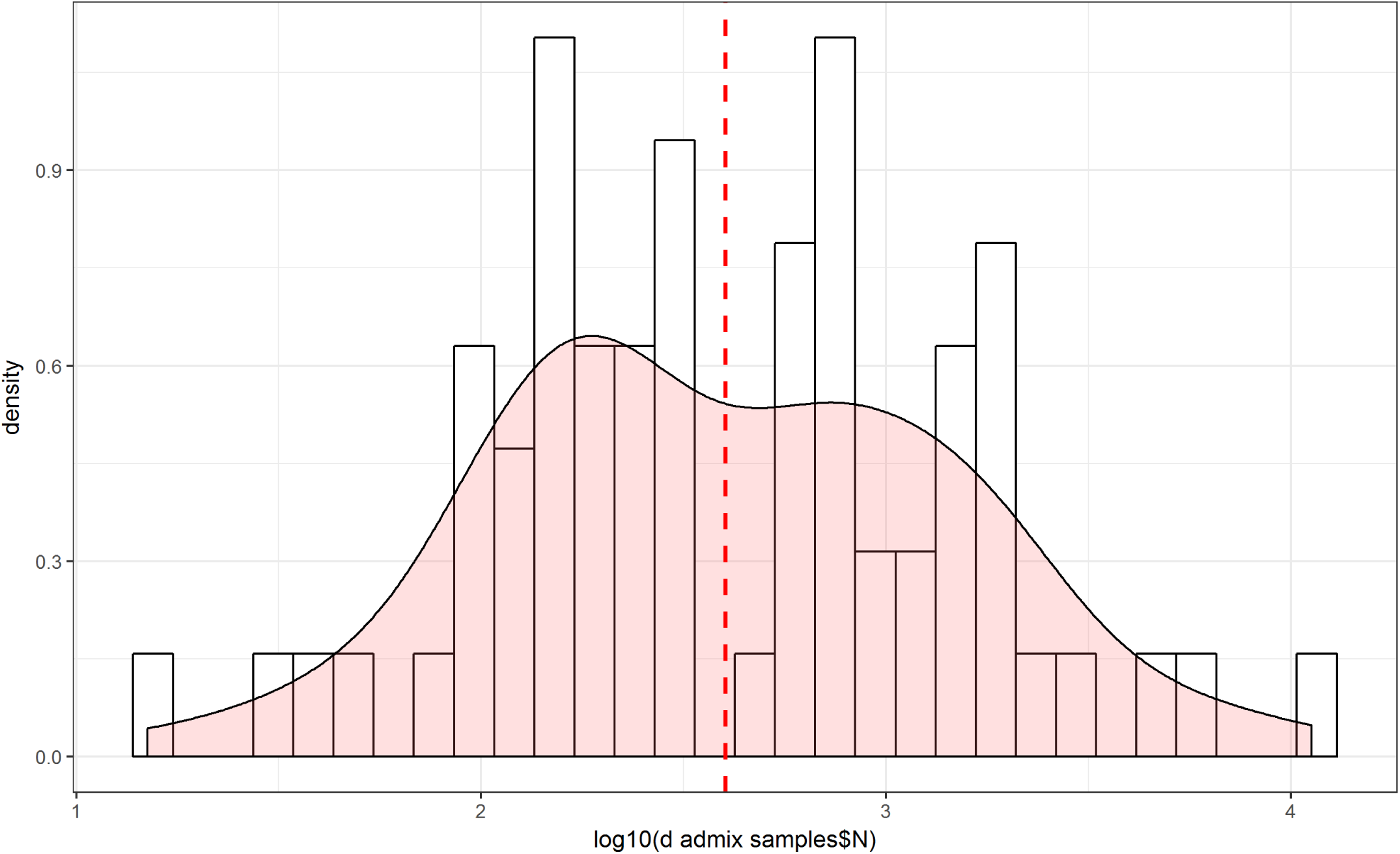
Density histogram of sample sizes, with a stippled line at the mean.

**Figure 3.**
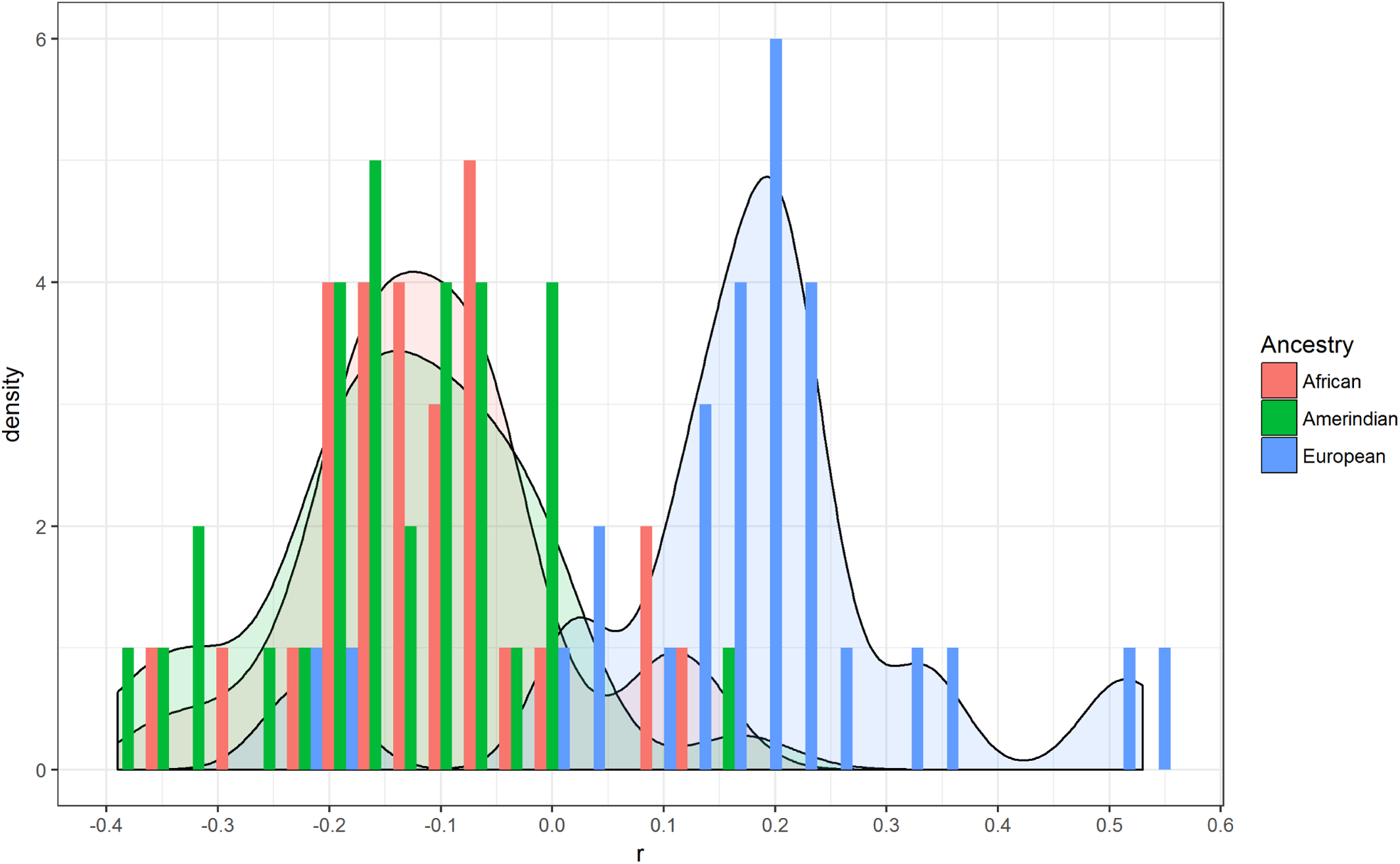
Density histogram plot for effect sizes by BGA.

**Table 6.** Descriptive statistics for the effect sizes

| BGA | N | K | Mean | Median | Max | Min | SD | 10<sup>th</sup><br>centile | 90<sup>th</sup><br>centile |
| --- | --- | --- | --- | --- | --- | --- | --- | --- | --- |
| African (w/o low admix) | 32710.5 | 28 | -.11 | -.12 | .14 | -.35 | .11 | -.21 | .03 |
| African (w low admix) | 37523.5 | 31 | -.09 | -.10 | .14 | -.35 | .11 | -.20 | .08 |
| Amerindian (w/o low admix) | 28937.5 | 31 | -.13 | -.13 | .17 | -.39 | .12 | -.31 | .00 |
| Amerindian (w low admix) | 40467.5 | 33 | -.13 | -.12 | .17 | -.39 | .12 | -.30 | .00 |
| European (w/o low admix) | 35476.5 | 28 | .17 | .19 | .53 | -.24 | .16 | .02 | .33 |
| European (w low admix) | 35476.5 | 28 | .17 | .19 | .53 | -.24 | .16 | .02 | .33 |

Note: $n$ = participant sample size; $K$ = number of independent samples. With/without BGA <5%.

There are two clear outliers for European BGA at -.24 and -.20. The first one was based on an extremely small sample (*n* = 15). The second one was both an outlier for European BGA (Zou et al. sample 6, *N* = 158) and for Amerindian BGA.^2^ When these outliers are excluded, the mean correlations changed little (*rs* = .16, -.14, and -.11 for European, Amerindian and African BGA respectively). Figures 4, 5 and 6 show the forest plots for the random effects models. Tables 7a to 7c show study details and effect sizes.

**Figure 4.**
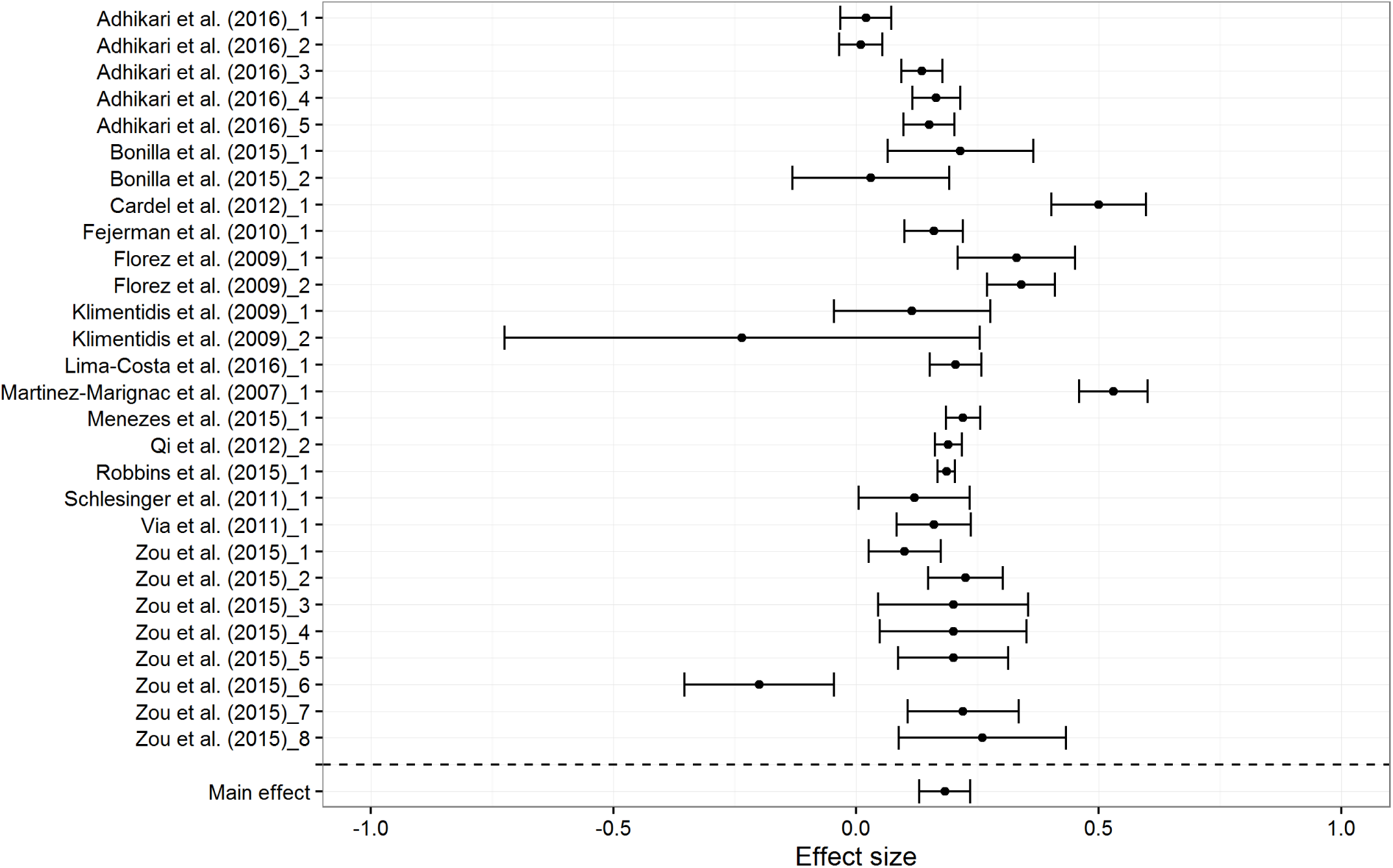
Forest plot for European BGA results, based on random effects model

**Figure 5.**
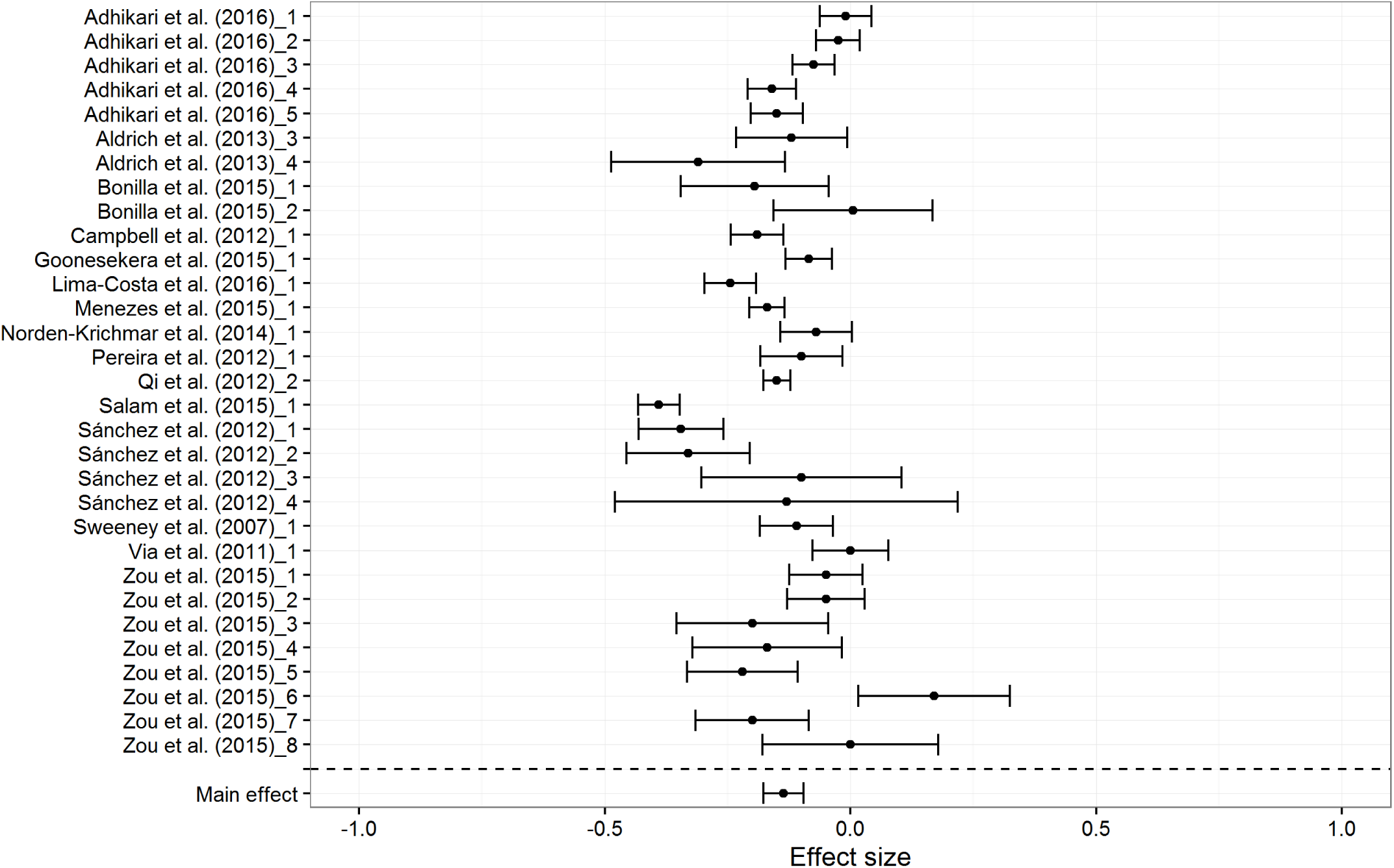
Forest plot for Amerindian BGA results, based on random effects model

**Figure 6.**
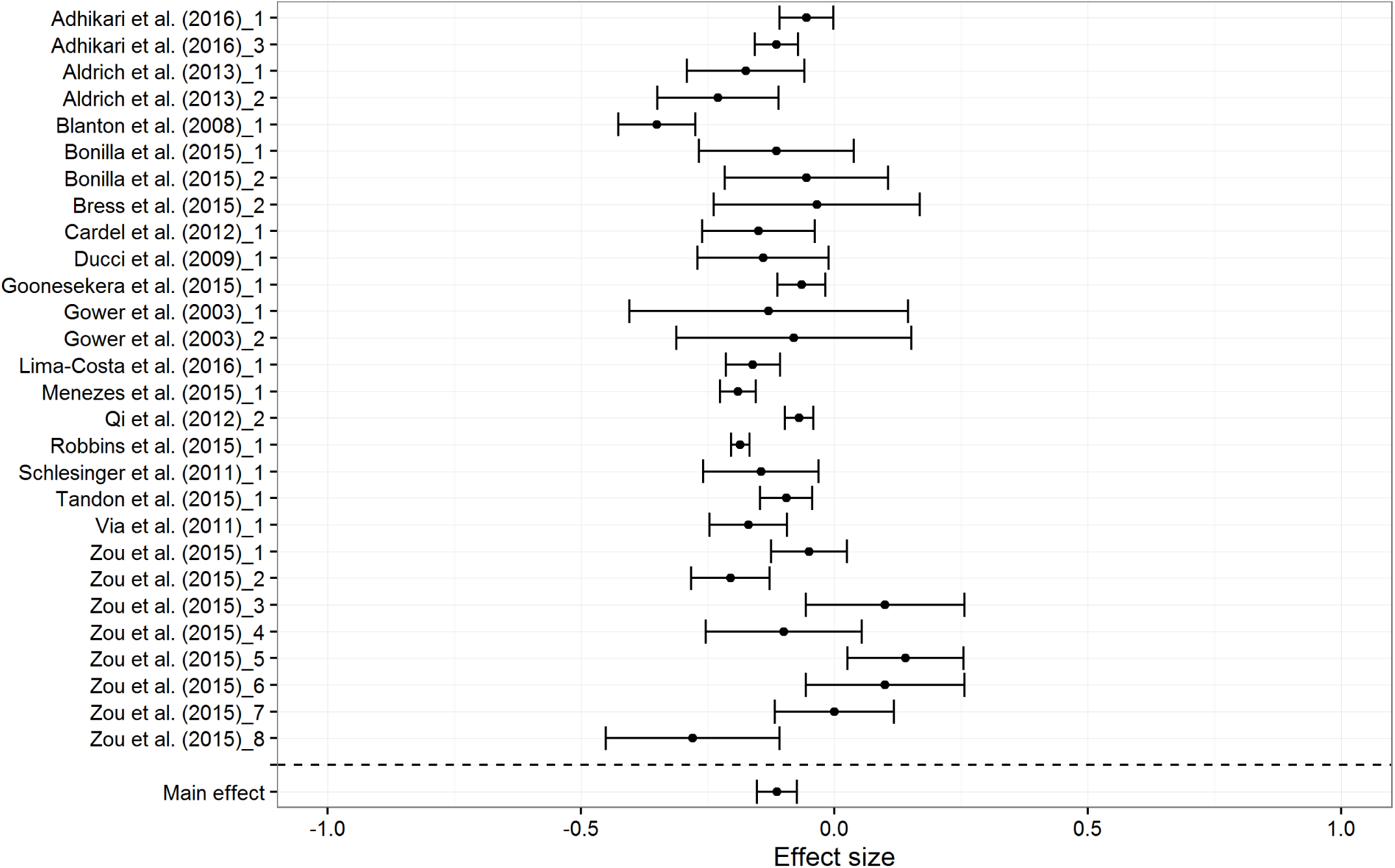
Forest plot for African BGA results, based on random effects model

**Table 7a.** Study details and effect sizes (with lower and upper 95% confidence intervals) for the meta-analyses: European BGA.

| Author sample | Country | Ethnic group | N | *r* | Lower | Upper |
| --- | --- | --- | --- | --- | --- | --- |
| Adhikari et al. (2016)_1 | Brazil | Multi-ethnic | 1375 | 0.02 | -0.03 | 0.07 |
| Adhikari et al. (2016)_2 | Chile | Multi-ethnic | 1934 | 0.01 | -0.03 | 0.05 |
| Adhikari et al. (2016)_3 | Colombia | Multi-ethnic | 2092 | 0.14 | 0.09 | 0.18 |
| Adhikari et al. (2016)_4 | Mexico | Multi-ethnic | 1535 | 0.16 | 0.12 | 0.21 |
| Adhikari et al. (2016)_5 | Peru | Multi-ethnic | 1344 | 0.15 | 0.10 | 0.20 |
| Bonilla et al. (2015)_1 | Uruguay | Not stated | 164 | 0.22 | 0.07 | 0.36 |
| Bonilla et al. (2015)_2 | Uruguay | Not stated | 148 | 0.03 | -0.13 | 0.19 |
| Cardel et al. (2012)_1 | US | Multi-ethnic | 305 | 0.50 | 0.40 | 0.60 |
| Fejerman et al. (2010)_1 | Mexico | Not stated | 1035 | 0.16 | 0.10 | 0.22 |
| Florez et al. (2009)_1 | Mexico | Not stated | 235 | 0.33 | 0.21 | 0.45 |
| Florez et al. (2009)_2 | Colombia | Not stated | 696 | 0.34 | 0.27 | 0.41 |
| Klimentidis et al. (2009)_1 | US | Hispanic | 147 | 0.12 | -0.05 | 0.28 |
| Klimentidis et al. (2009)_2 | US | Native American | 15 | -0.24 | -0.72 | 0.25 |
| Lima-Costa et al. (2016)_1 | Brazil | Not stated | 1305.5 | 0.20 | 0.15 | 0.26 |
| Martinez-Marignac et al. (2007)_1 | Mexico | Not stated | 561 | 0.53 | 0.46 | 0.60 |
| Menezes et al. (2015)_1 | Brazil | Multi-ethnic | 2960 | 0.22 | 0.18 | 0.26 |
| Qi et al. (2012)_2 | US | Hispanic | 4980 | 0.19 | 0.16 | 0.22 |
| Robbins et al. (2015)_1 | US | African-American | 11239 | 0.19 | 0.17 | 0.20 |
| Schlesinger et al. (2011)_1 | Brazil | Not stated | 291 | 0.12 | 0.01 | 0.23 |
| Via et al. (2011)_1 | US | Puerto-Rican | 642 | 0.16 | 0.08 | 0.24 |
| Zou et al. (2015)_1 | US | Puerto-Rican | 686 | 0.10 | 0.03 | 0.17 |
| Zou et al. (2015)_2 | US | Puerto-Rican | 615 | 0.23 | 0.15 | 0.30 |
| Zou et al. (2015)_3 | US | Mexican | 157 | 0.20 | 0.05 | 0.35 |
| Zou et al. (2015)_4 | US | Mexican | 163 | 0.20 | 0.05 | 0.35 |
| Zou et al. (2015)_5 | US | Mexican | 291 | 0.20 | 0.09 | 0.31 |
| Zou et al. (2015)_6 | US | Mexican | 158 | -0.20 | -0.35 | -0.05 |
| Zou et al. (2015)_7 | US | Mexican | 281 | 0.22 | 0.11 | 0.33 |
| Zou et al. (2015)_8 | US | Puerto-Rican | 122 | 0.26 | 0.09 | 0.43 |
| **RE Model** |  |  | **35476.5** | **0.18** | **0.13** | **0.24** |

**Table 7b.**
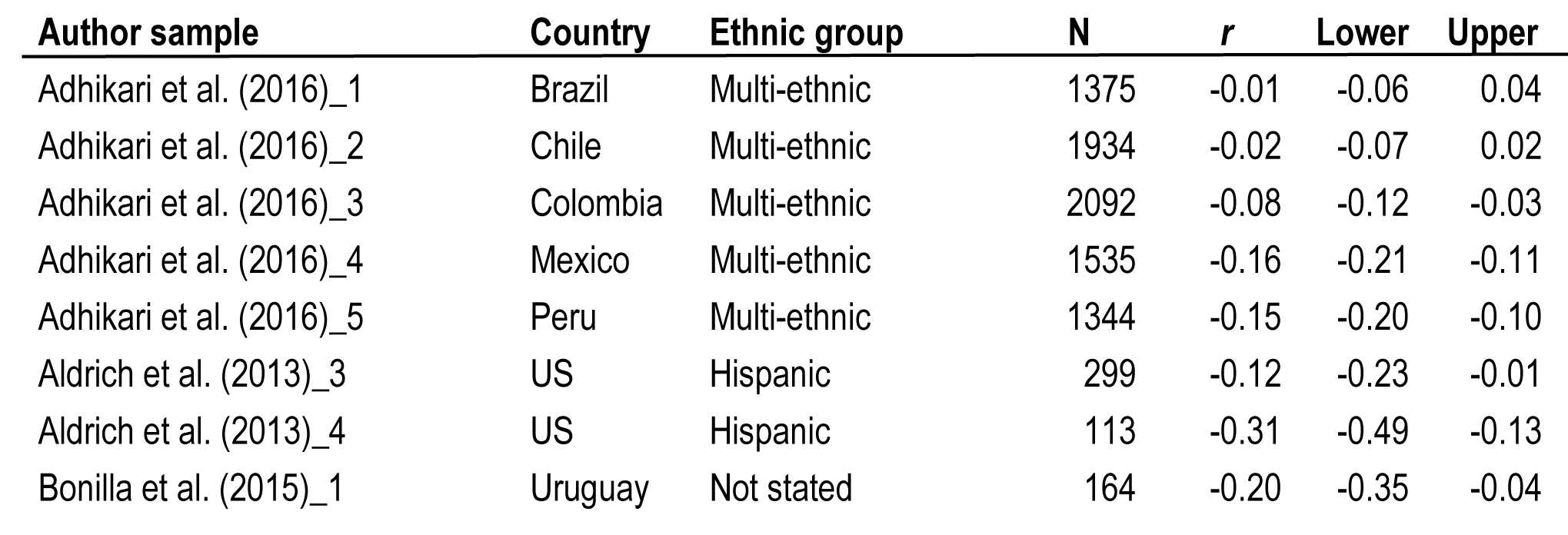
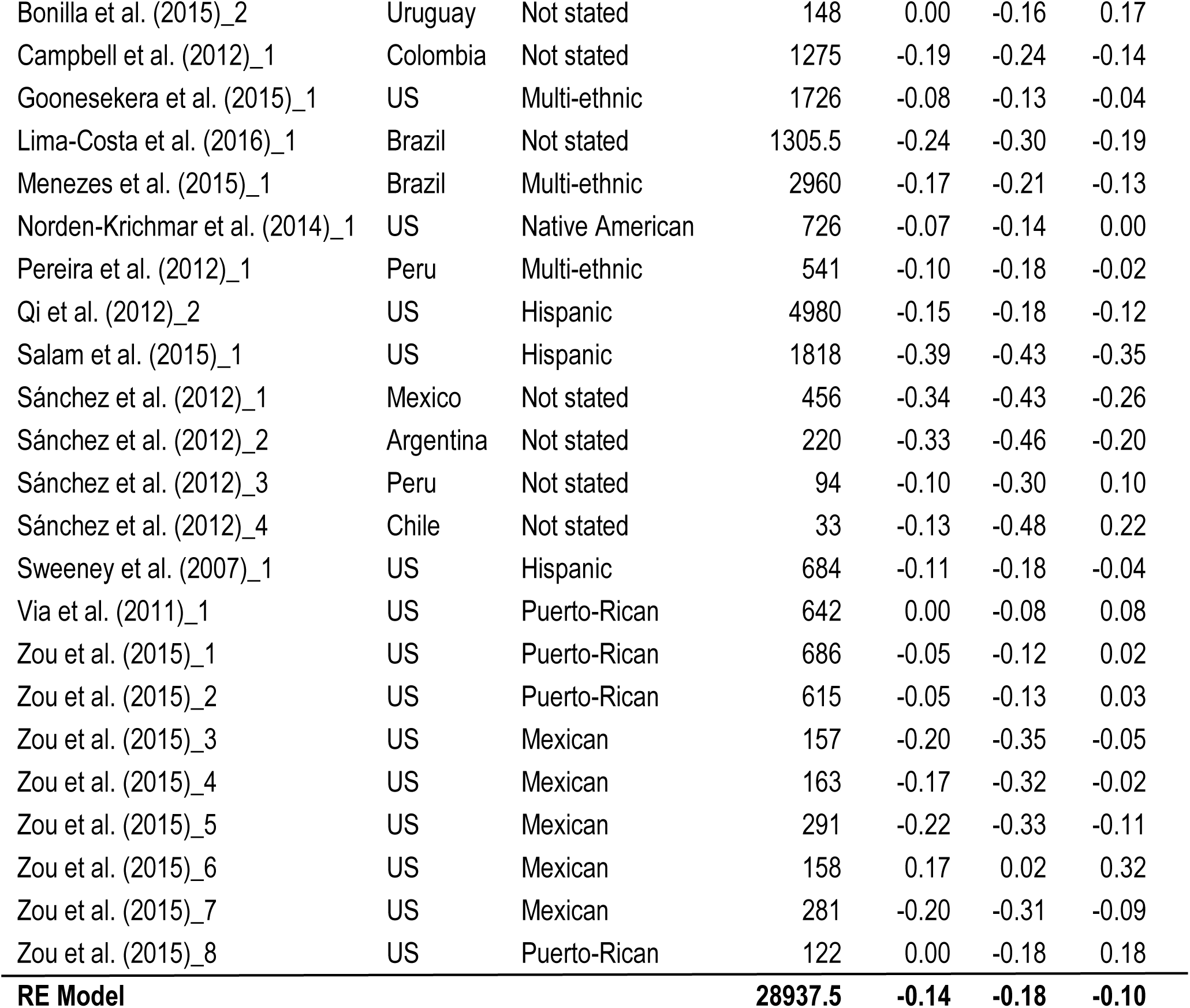
Study details and effect sizes (with lower and upper 95% confidence intervals) for the meta-analyses: Amerindian BGA.

**Table 7c.**
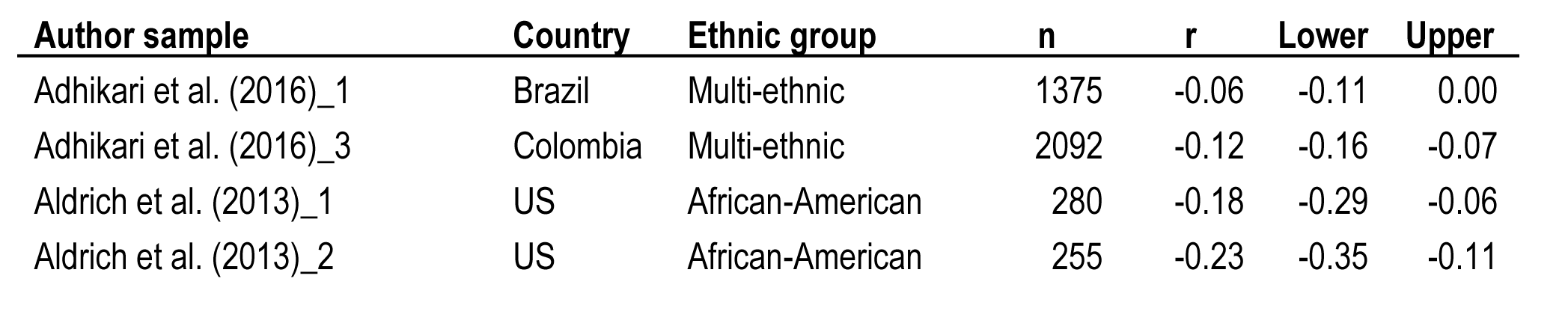
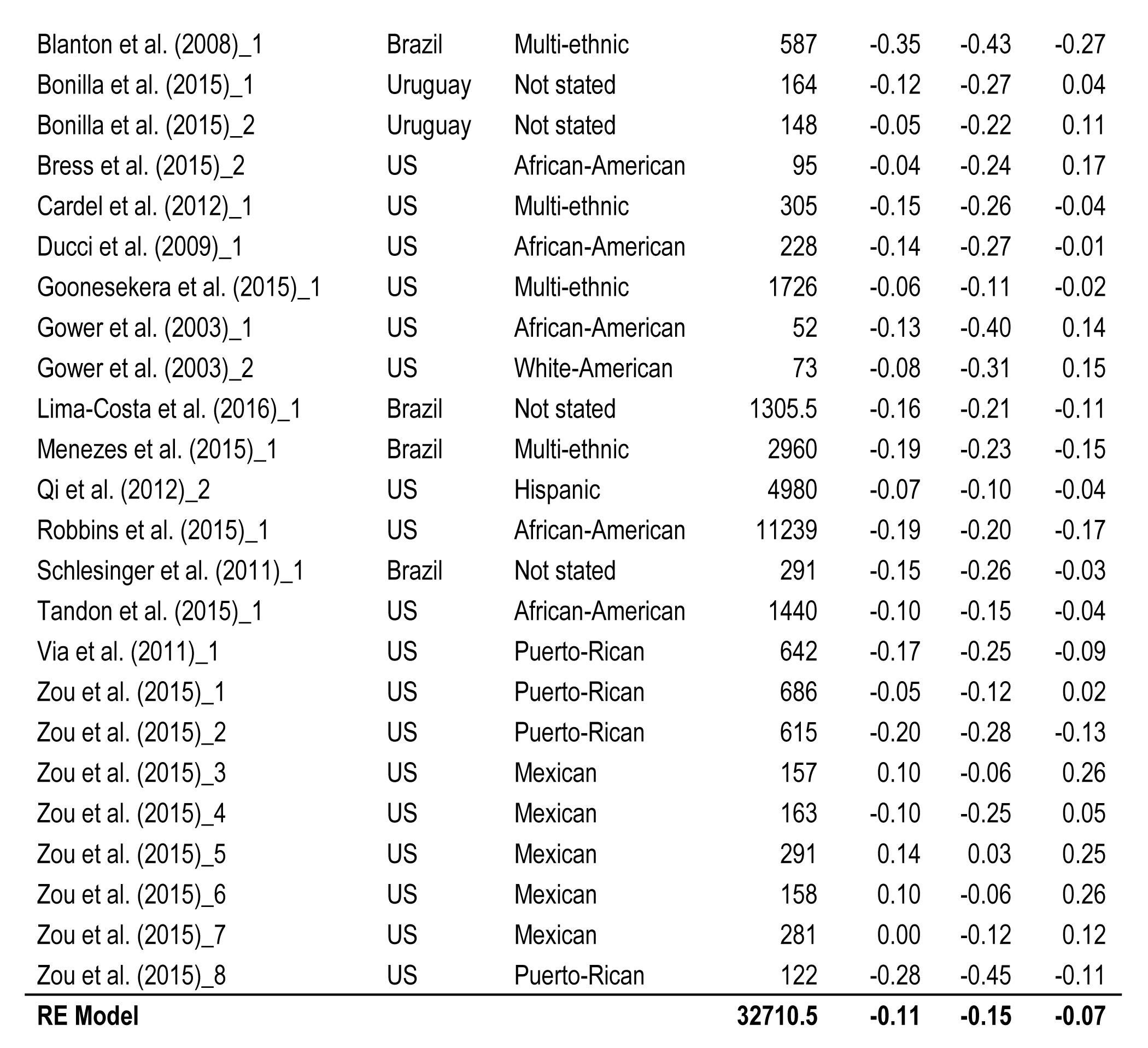
Study details and effect sizes (with lower and upper 95% confidence intervals) for the meta-analyses: African BGA.

The random effect meta-analytic mean effect sizes are .18, -.14, and -.11 for European, Amerindian and African BGA, respectively. European BGA shows a stronger association with SES than do Amerindian and African BGA, presumably because it is the largest ancestry component (mean European BGA = 54.41%). The difference could be due to the higher variation in the proportion of European BGA in the subjects. The mean standard deviation of admixture (18.01, 11.18, and 14.50 for European, Amerindian and African, respectively) was lower for the non-European ancestries, and less variation is expected to lead to smaller effect sizes owing to restriction of range (Schmidt & Hunter, 2014).

The exclusion of samples with a mean BGA below 5% had only a slight impact on the results. When all samples are included, the meta-analytic effect sizes are *r* = .18 [.13, .24], *r* = -.13 [-.17, -.09], and r = -.09 [-.13, -.05] for European, Amerindian and African BGA, respectively. Results for African BGA are impacted the most owing to the presence of a positive association between African BGA and outcomes for several samples with very low African admixture (e.g., Adhikari et al., 2016: Chile, African admixture 2.61 %, *r* BGA ⨯ SES = .08). It is notable that for two of the same countries, namely Chile and Mexico, Fuerst and Kirkegaard (2016a, 2016b) found that regional African BGA is associated with higher SES.

The funnel plots did not show notable asymmetry, suggesting no publication bias. Figures 7, 8, and 9 show, respectively, the funnel plots for European, Amerindian, and African BGA.

**Figure 7.**
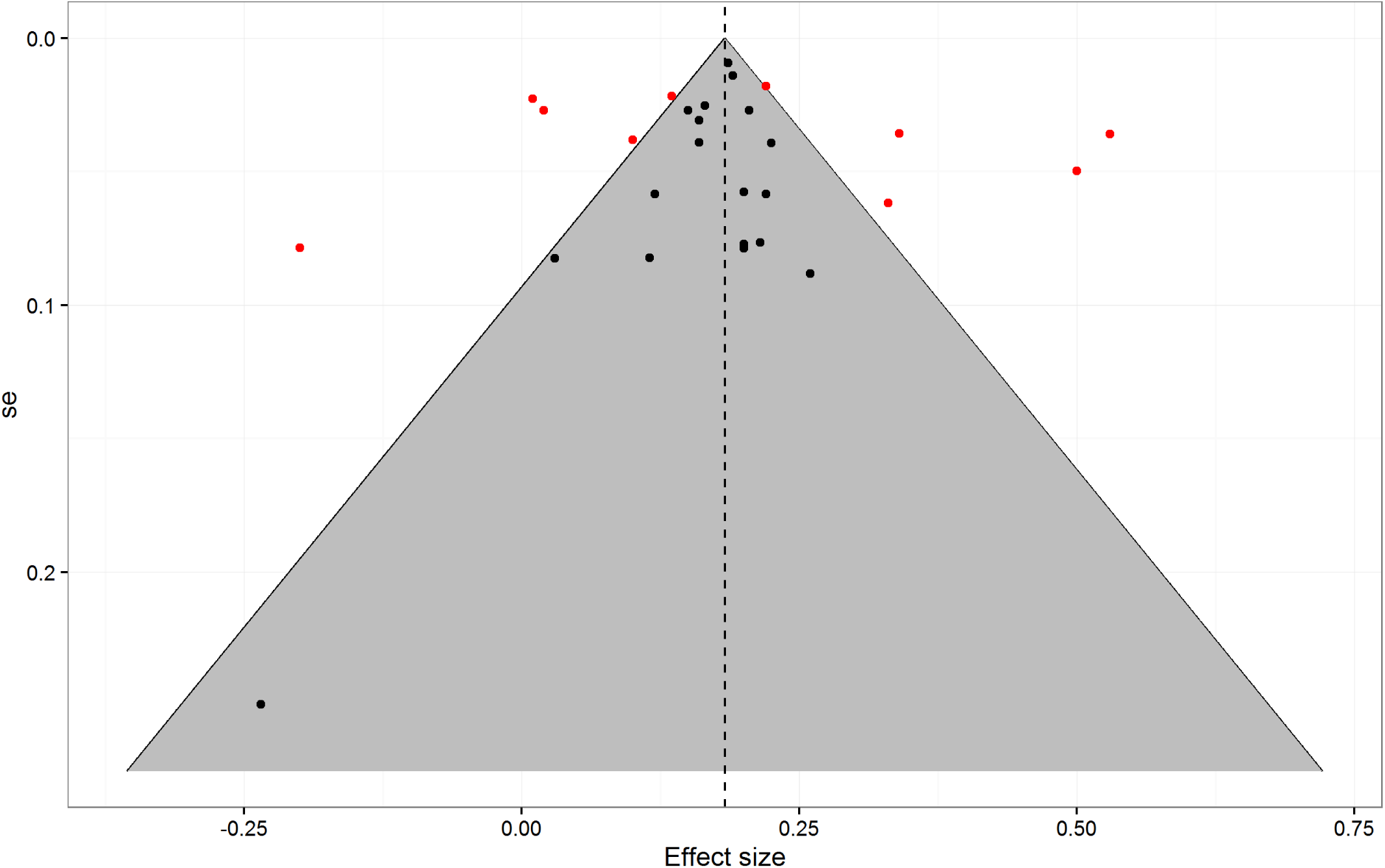
Funnel plot for European BGA, with standard error on the y-axis and effect size on the x-axis

**Figure 8.**
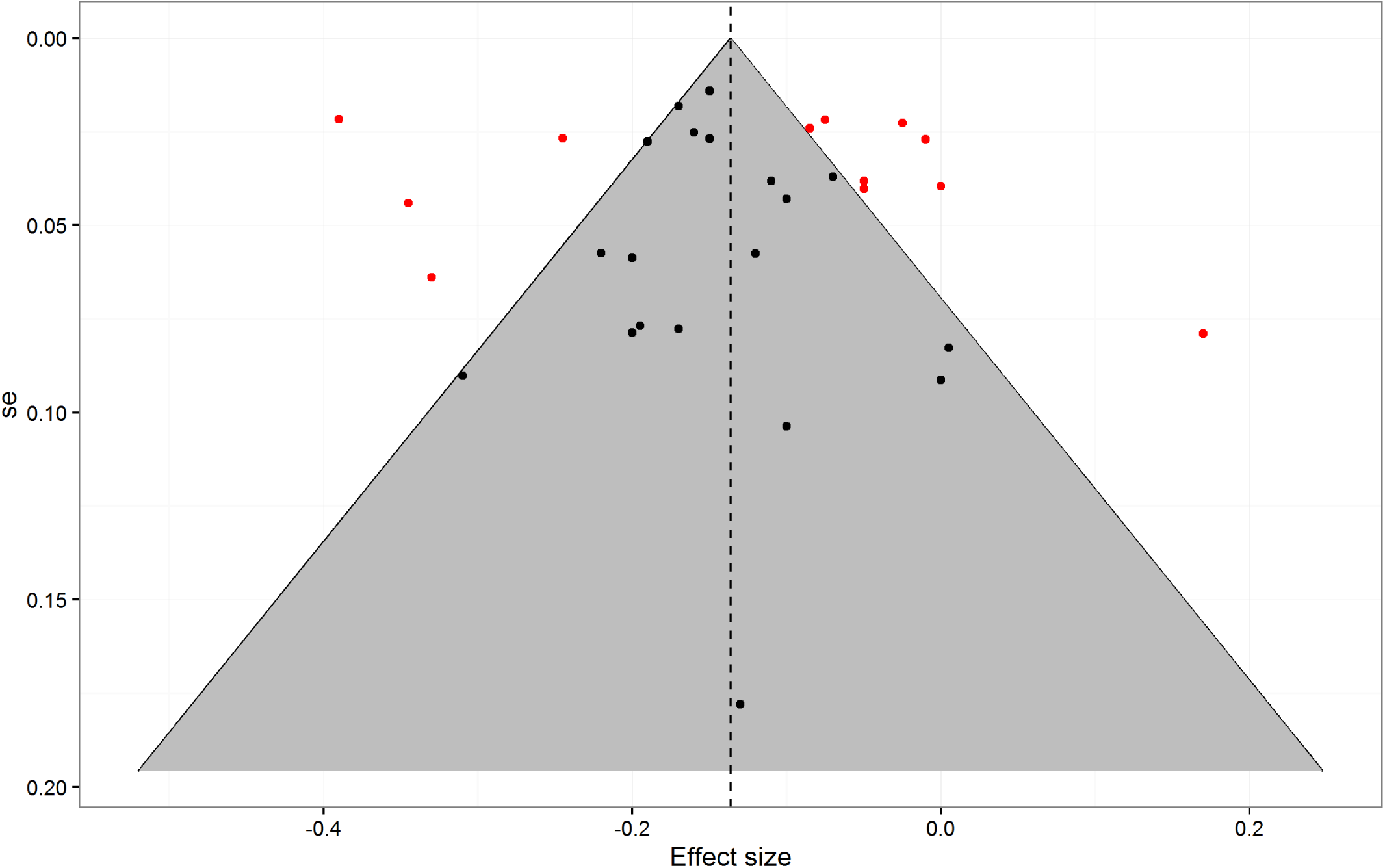
Funnel plot for Amerindian BGA, with standard error on the y-axis and effect size on the x-axis

**Figure 9.**
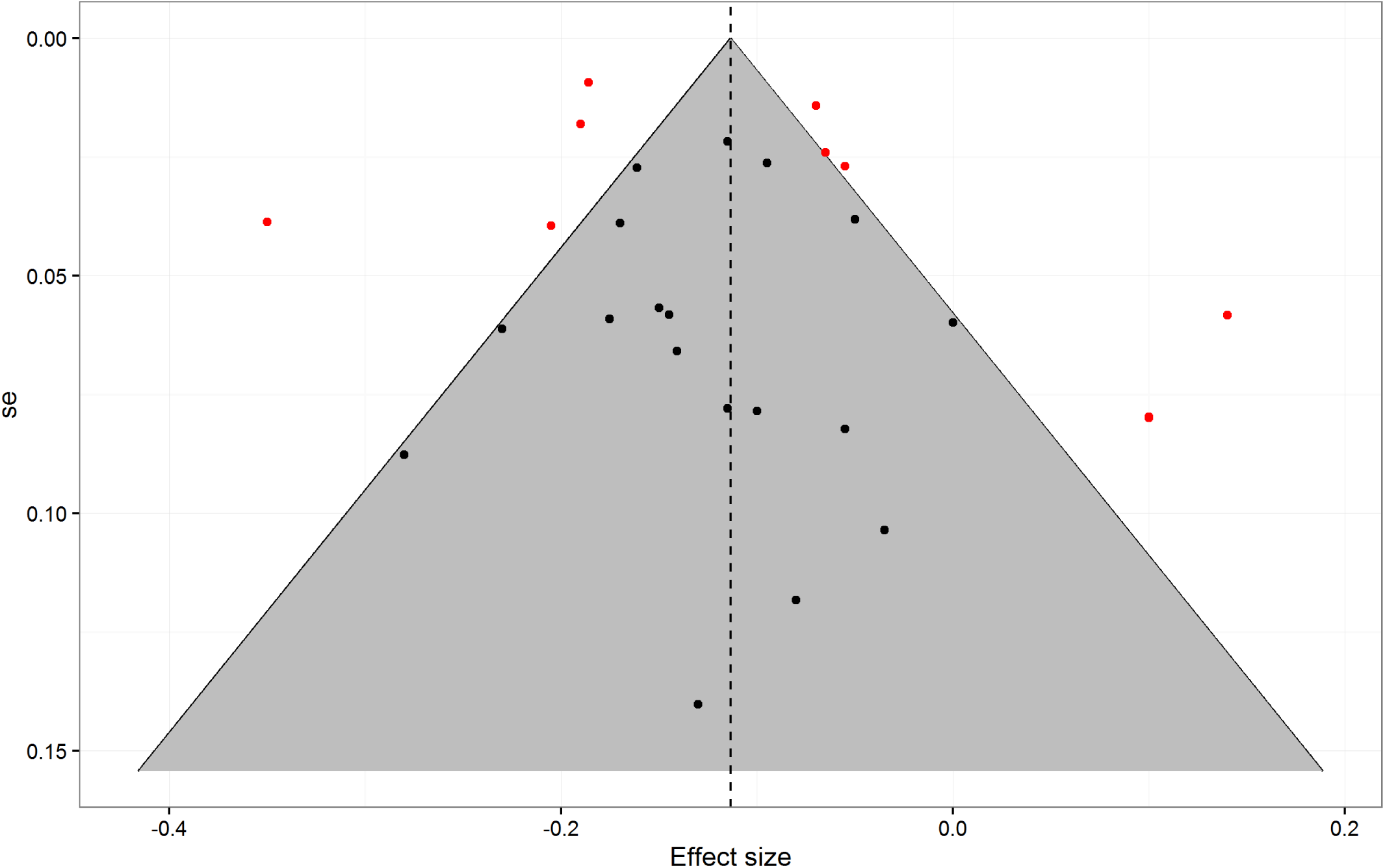
Funnel plot for African BGA, with standard error on the y-axis and effect size on the x-axis

## 3. Discussion

### 3.1. Main results

The results of the meta-analysis are consistent with those reported in earlier narrative reviews in that European BGA was statistically associated with more favorable socioeconomic outcomes relative to Amerindian and African BGA.

When low-admixture samples were excluded, the random effects meta-analysis gave a mean SES ⨯ BGA correlation of *r* = .18, *r* = -.14, and *r* = -.11 for European, Amerindian and African BGA, respectively; furthermore, the analysis of SES ⨯ BGA association directions gave square root *n*-weighted means of *M* = 0.83, *M* = -0.81, and *M* = -0.82 for European, Amerindian and African BGA, respectively. For the random effects meta-analysis, the heterogeneity, or the percentage of variance due to variance between samples as opposed to sampling error, was considerable (I^2^ _mean_ = 92%, I^2^ _European_ = 95%, I^2^ _Amerindian_ = 91%, I^2^ _African_ = 89%) by conventional standards (Higgins & Green, 2008). Due to the limited number of samples, it is difficult to evaluate the cause of this pattern of results. Some possibilities are as follows:

1.*Discretization:* Many outcome variables were ordinal instead of continuous, even when continuous values were available (e.g., income). Pearson correlations assume that the data are normally and continuously distributed, so the use of non-continuous data induces a downward bias in the results. Corrections for discretization were not attempted, although one could attempt to apply these (Schmidt & Hunter, 2014).

2.*Number of BGA-informative markers:* There were large differences in the number of genomic markers used to estimate individual BGA. Figure 10 shows a density histogram of the distribution for the effect size analysis with low-admixture samples removed. Using fewer markers results in more measurement error with respect to true BGA (Russo et al., 2016). Ruiz-Linares et al. (2014) reported correlations between admixture estimates using different numbers of markers. They noted that a recent study (Scharf et al., 2013) had found that using 15 markers resulted in correlations of about .60 with estimates derived from 50k markers. Furthermore, using 30 and 152 markers resulted in *r* = .70 and *r* = .85, respectively, with respect to estimated admixture based on 50k markers. It is clear that there are diminishing returns to using more markers, but also that using more reduces measurement error with respect to true BGA. It would be difficult to correct for measurement error with respect to BGA. One option would be to acquire a sufficient number of data points from one study to allow for the estimation of a predictive model. One could then use that model to estimate the measurement error in other studies based on the number of reported markers used. However, we failed to find a study which had sufficient data points, so a correction was not attempted.

**Figure 10.**
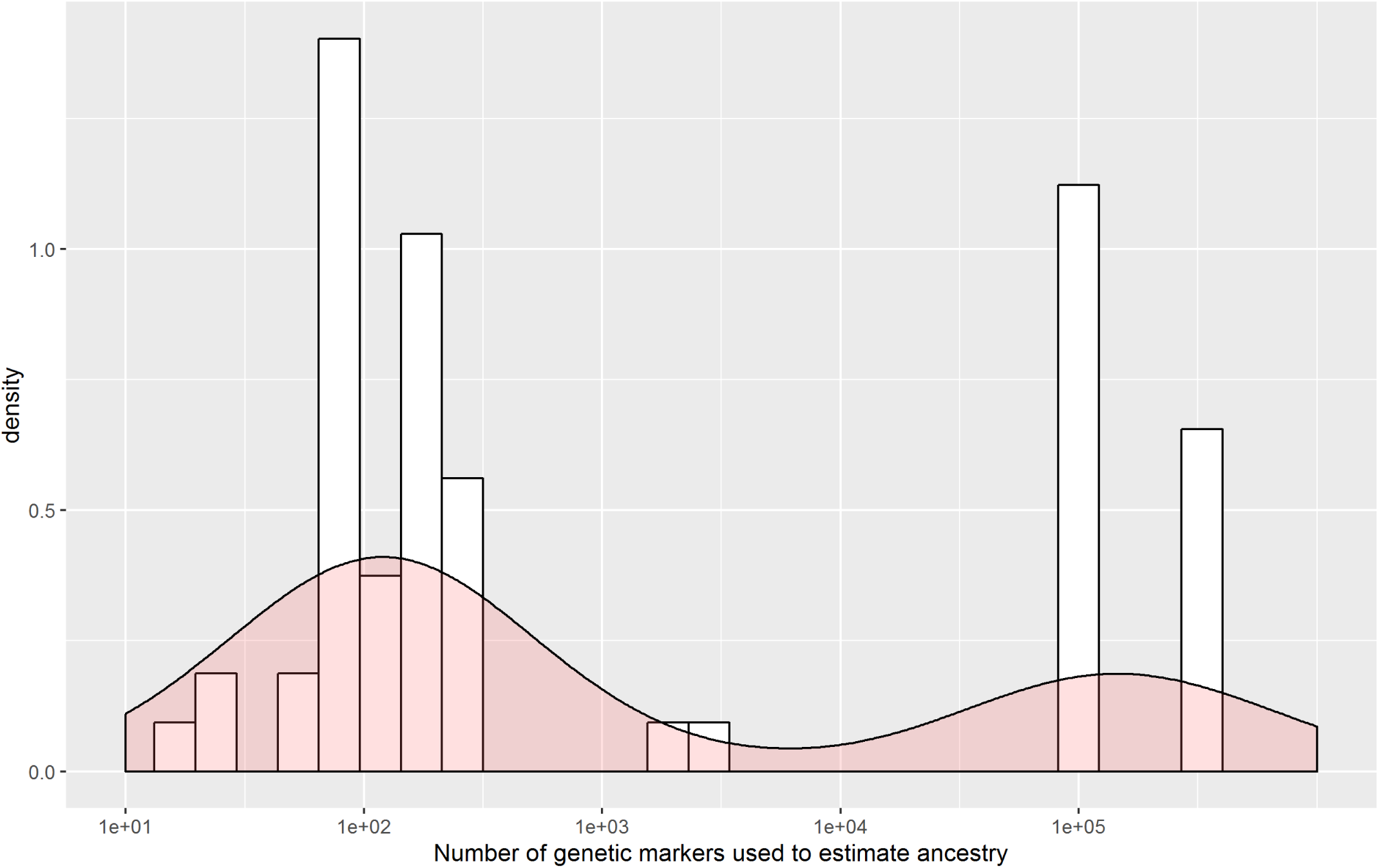
Density histogram of the number of genetic markers in each sample. Average frequency density on the y-axis and number of genetic markers on the x-axis.

3. *Variation in admixture:* Based on the samples for which standard deviations of BGA were reported, variability in BGA differed considerably between samples. For instance, for Uruguayans, Bonilla et al. (2015) found a standard deviation of African BGA of 7.52; in contrast, for Brazilians, Schlesinger et al. (2011) found one of 24.08. As mentioned earlier, the magnitude of associations between outcomes and BGA will be impacted by variance in BGA. Thus, between-sample heterogeneity in BGA variance is expected to contribute to heterogeneity in BGA ⨯ SES associations.

4. *Heterogeneous origin:* The studies in the meta-analysis came from many countries. It is probable that the statistical effect of one’s BGA depends on local cultural norms or practices that differ between countries or even between regions within countries.

5. *Sampling:* Most of the studies used convenience samples, an approach that can have an impact on associations. Adhikari et al.’s (2016) samples, for example, were recruited mostly at universities, meaning that individuals were selected with respect to educational status. Thus, the correlations they obtained are expected to be attenuated relative to those derived from more representative samples.

### 3.2. Implications for epidemiological studies

Fairly robust associations between BGA and SES were found. Given this, to avoid spurious associations in regression analyses due to omitted variable bias, it is potentially important to include reliable measures of SES in studies of medical outcomes. Studies reviewed in this analysis employed a wide variety of specific measures (see Table 3), some of which may be better indices of medically relevant SES than others. It would be worthwhile to investigate which measures (neighborhood SES, education, income, etc.) are generally associated to a greater degree with BGA and with medical outcomes, so that future studies could attempt to incorporate these measures into analyses.

Further, it is also important to identify the factors mediating the BGA-SES associations, as these could incrementally explain the BGA-medical outcome associations. Figure 11 shows one possible model. In this model, local ancestry is a subset of global ancestry. Both local and global ancestry correlate with disease loci which, in turn, are causally related to health outcomes. Global ancestry is also related both to an individual’s SIRE and to cultural practices associated with SIRE, and to an individual’s race/ethnicity-associated phenotype, and social reactions, perhaps in the form of “colorism” (e.g., Hunter, 2007; Telles, 2014), associated with that phenotype. In this model, both direct and indirect pathways run from cultural practices and social reactions to health outcomes. An example of a direct pathway would be through observed race/ethnicity-related differential treatments by providers (as discussed by, e.g., Van Ryn, 2002). Indirectly, these factors potentially impact both individuals’ human capital and SES, which are reciprocally related to one another and to health outcomes.

**Figure 11.**
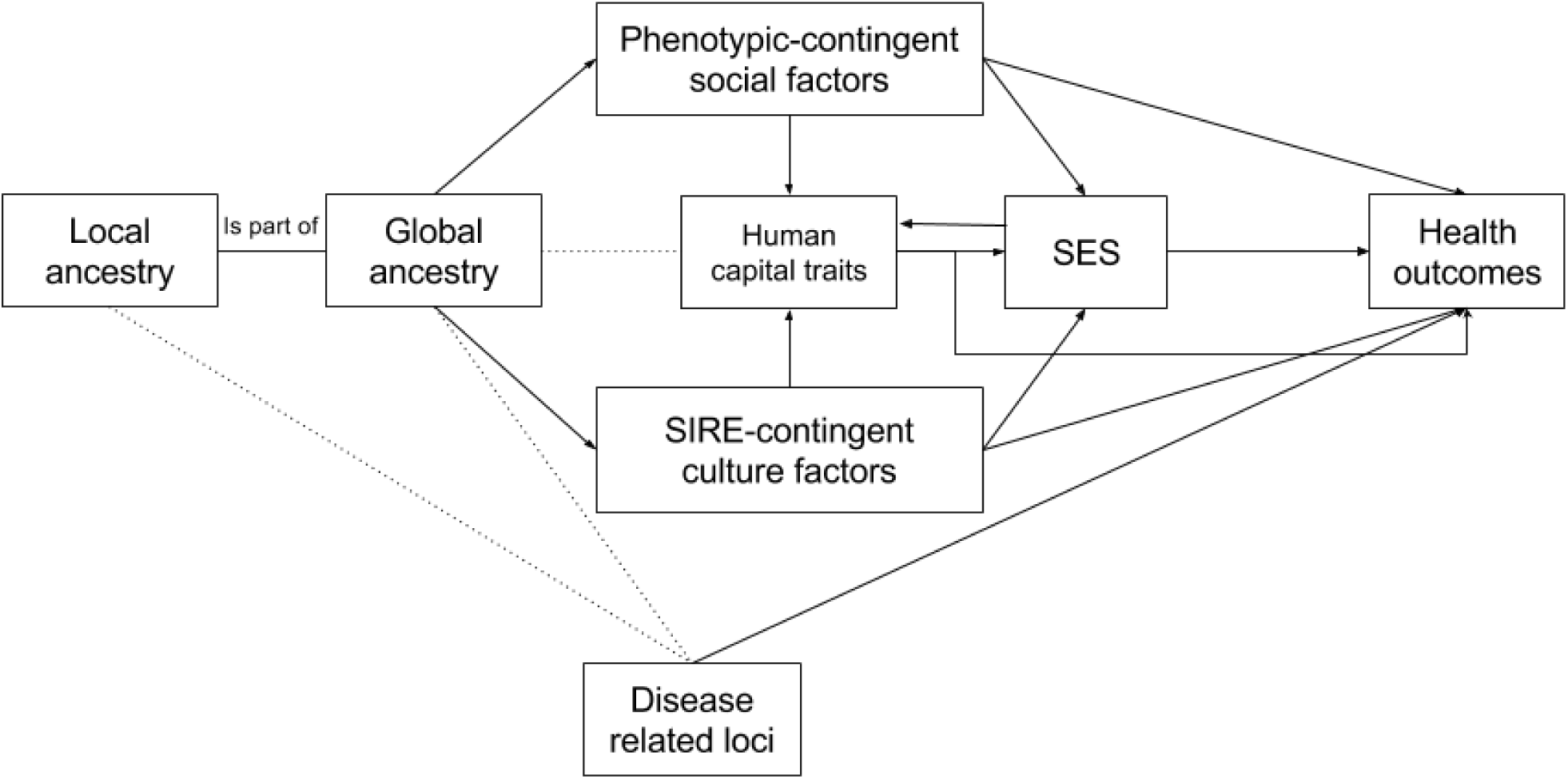
Model of possible relations between BGA and SES. The stippled lines represent correlations and the arrowed lines represent causal associations.

This model is similar to that proposed by Marden et al. (2016). A notable exception is that we include human capital as a mediating variable. Cognitive epidemiology studies have shown that measures of cognitive ability (one component of human capital) can explain a portion of medical-related outcomes, independent of SES (Deary, 2009; Der, Batty & Deary, 2009; Wraw et al., 2015). Because, throughout the Americas, SIRE groups are known to differ in mean levels of phenotypic cognitive ability (Fuerst & Kirkegaard, 2016b; Roth et al., 2001), it would be reasonable to include measures of cognitive ability in admixture analyses to see if such measures explain some of the health outcome differences independent of SES. However, we could not find any study that did this. On the other hand, we did locate one recent admixture paper (Akshoomoff et al., 2014) which reported that BGA was associated with cognitive ability. On inquiry, as expected given the SIRE differences, European BGA was positively associated with cognitive ability, while African and Amerindian BGA were negatively associated (Akshoomoff, personal communications, November 9, 2014). If cognitive ability mediates the relationship between BGA and SES, not including measures of it constitutes omitted variable bias.

### 3.3. Untangling the statistical effects of BGA and SIRE

In many of the samples included in this meta-analysis, individual SES outcomes are associated with BGA within SIRE groups. Hence, SIRE membership does not mediate the relationship. In other cases, particularly in Latin America, the issue is less clear and BGA may be confounded with SIRE. For instance, Leite et al. (2011) found that European BGA was positively correlated with SES in Brasília. This could be because SES was positively correlated with BGA independently of SIRE, or because it was positively correlated with SIRE but not with BGA independent of SIRE. The analysis by Ruiz-Linares et al. (2014) has helped to clarify the issue. The authors looked at the association between BGA, skin pigmentation and SIRE in a multi-country sample from Brazil, Chile, Colombia, Mexico and Peru (mean age 20 to 25, depending on country). The authors found modest correlations between BGA and SIRE (e.g., 0.48 in the case of both European and Amerindian BGA/SIRE). They reported that wealth and educational attainment correlated with European BGA (r = .12 for the full sample). However, when the statistical effect of BGA was controlled for, education was not significantly associated with SIRE, and wealth was only related to SIRE within the European/White color group. Similar results in this regard were found with Menezes et al.’s (2015) sample from the 1982 Pelotas Birth Cohort study (F. Hartwig, personal communication, March 4, 2016). Details pertaining to this sample can be found in Menezes et al. (2015). The results pertaining to self-identified color and interviewer-rated color are shown in Supplementary File 3. For this sample, the positive association between European BGA and household assets, schooling, and income was robust to controls for interviewer-observed race/ethnicity and SIRE. Contrary to a simple colorism model, Black (*preta*) and Brown (*parda*) interviewer-rated race/ethnicity were generally positively related to outcomes, controlling for European BGA. Taken together, the results suggest that the association between BGA and SES is substantially independent of SIRE and thus SIRE-related cultural factors. Quite possibly, phenotype-based social reactions, which would differentially impact individuals within SIRE groups, affect outcomes. For example, consistent with this possibility, Telles (2014) found, in a large multinational sample, that interviewer-rated skin brightness was related to SES controlling for SIRE.

### 3.4. Limitations and suggestions for future research

The number of effect sizes used in the present meta-analysis is limited. This is because many studies did not report effect sizes and, in some cases, the authors either did not reply to emails or were unable to provide results or data. Meta-analyses would be more reliable and easier to perform if scientists were more willing to publish their data and report effect sizes. The results significantly varied across studies (mean effect size heterogeneity = 92%), which means that there are effect size moderators. A moderator analysis was not conducted due to the relatively small number of studies in the dataset. We suggest that moderator analyses be conducted as relevant data accumulates. Detailed analyses for publication bias were also not conducted because the number of effect sizes was too small for a reliable analysis. This concern should be investigated in future meta-analyses.

## Supplementary material

Supplementary File 1-3, high-quality figures and R analysis code are available at the repository at *Open Science Framework* https://osf.io/ydc3f/files/.

*indicates that the study included data used in the meta-analysis.

## Footnotes

2 The lead author (James Zou) was contacted to verify the data. He checked the code and data and reported that there were no errors.

